# DeepSLICEM: Clustering CryoEM particles using deep image and similarity graph representations

**DOI:** 10.1101/2024.02.04.578778

**Authors:** Meghana V. Palukuri, Edward M. Marcotte

## Abstract

Finding the 3D structure of proteins and their complexes has several applications, such as developing vaccines that target viral proteins effectively. Methods such as cryogenic electron microscopy (cryo-EM) have improved in their ability to capture high-resolution images, and when applied to a purified sample containing copies of a macromolecule, they can be used to produce a high-quality snapshot of different 2D orientations of the macromolecule, which can be combined to reconstruct its 3D structure. Instead of purifying a sample so that it contains only one macromolecule, a process that can be difficult, time-consuming, and expensive, a cell sample containing multiple particles can be photographed directly and separated into its constituent particles using computational methods.

Previous work, SLICEM, has separated 2D projection images of different particles into their respective groups using 2 methods, clustering a graph with edges weighted by pairwise similarities of common lines of the 2D projections. In this work, we develop DeepSLICEM, a pipeline that clusters rich representations of 2D projections, obtained by combining graphical features from a similarity graph based on common lines, with additional image features extracted from a convolutional neural network. DeepSLICEM explores 6 pretrained convolutional neural networks and one supervised Siamese CNN for image representation, 10 pretrained deep graph neural networks for similarity graph node representations, and 4 methods for clustering, along with 8 methods for directly clustering the similarity graph. On 6 synthetic and experimental datasets, the DeepSLICEM pipeline finds 92 method combinations achieving better clustering accuracy than previous methods from SLICEM. Thus, in this paper, we demonstrate that deep neural networks have great potential for accurately separating mixtures of 2D projections of different macromolecules computationally.

## Introduction

Awarded the Nobel Prize in Chemistry in 2017, cryogenic-electron microscopy (cryo-EM), a method for imaging flash-frozen samples of macromolecules [1] with electron beams, has produced more than 32k structures of various macromolecules in the Electron Microscopy Databank (EMDB), many of near-atomic resolution, as of February 2024. While growth in the number of structures solved by cryo-EM has been exponential [2], most structures have been determined from studies using single-particle cryo-EM on highly purified samples, although cryo-electron tomographic (cryo-ET) methods are increasingly revealing structures in cellular contexts (e.g., [3]). Importantly, determining the structures of macromolecules has several applications, as for instance, the cryo-EM structure of the SARS-CoV-2 spike protein [4] helped develop vaccines for COVID-19.

Recently, the cryo-EM technique has been extended to identifying the most abundant particles in more complex cellular extracts [5], [6] rather than in homogenous purified samples. This approach presents the additional challenge of computationally sorting images from different macromolecules prior to reconstructing their 3D structures. Despite this challenge, in principle, many more structures can be solved by computationally separating different particles from micrographs of heterogeneous mixtures of macromolecules in a cellular extract (the process is visualized in **Figure 1**).

**Figure 1.**
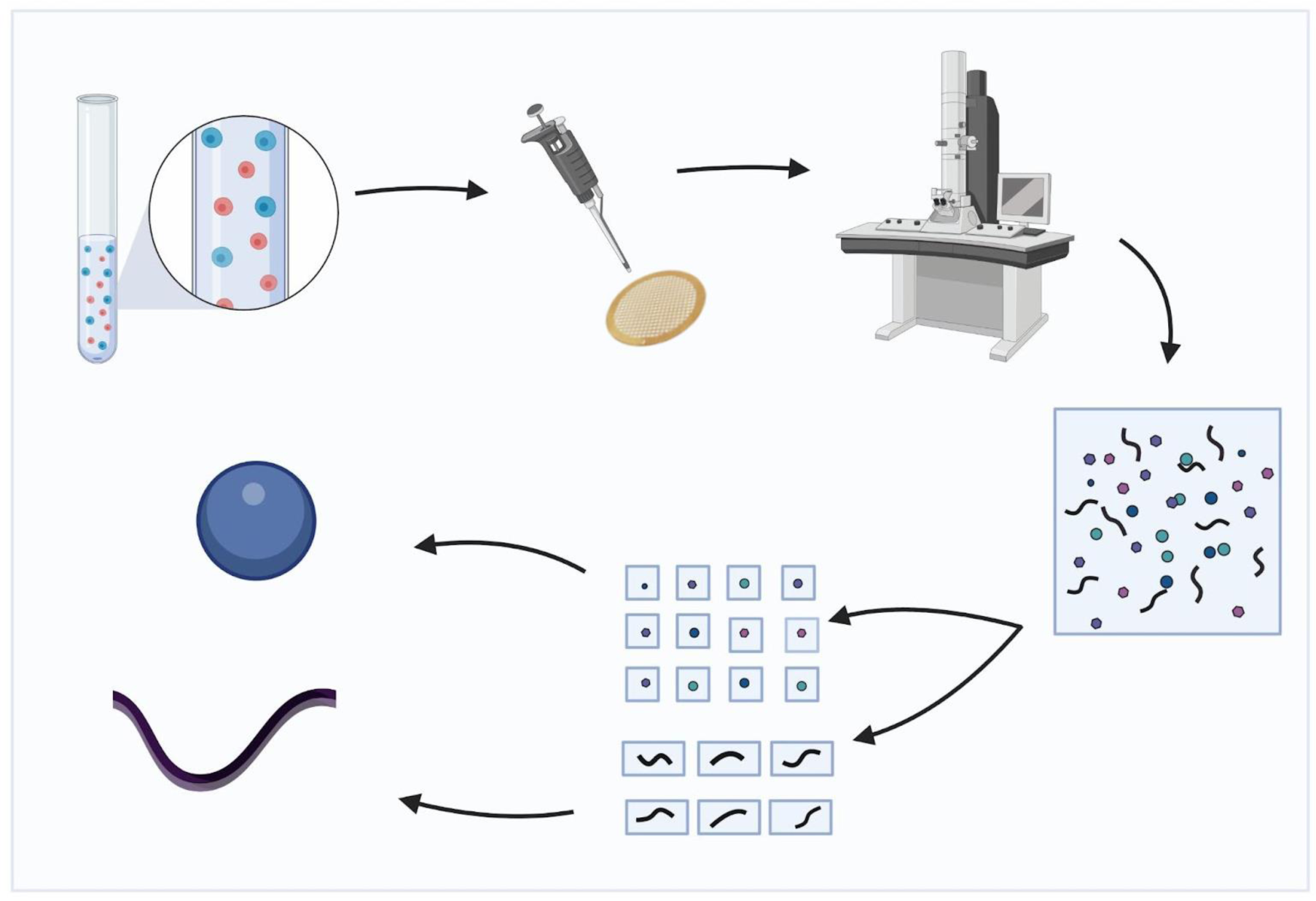
Overview of the cryo-EM structure determination process for multiple particles in a mixture. A sample with multiple macromolecules is photographed with a cryo-electron microscope. From the resulting micrograph, 2D projections of constituent particles are selected and separated into different groups corresponding to distinct particles. Each group of 2D projections is used to reconstruct each of the 3D structures of the different particles.

Previous methods for analyzing such data compute similarities between projections of particles photographed and cluster the projections into groups corresponding to particles; however, most of these methods [7]–[9] have been demonstrated to solve a similar problem of separating heterogeneous mixtures of different conformations of the same particle. When applied to heterogeneous mixtures of different particles, SLICEM [10] takes advantage of the property of the projection slice theorem in cryo-EM [11]. This theorem states that a common line, *i.e.*, a common 1D line projection, exists for any two 2D projections of the same 3D object. Thus, SLICEM uses common lines of 2D projections, constructing a graph with the similarity of the common lines of two projections as the edge weight and finally clusters the graph to group 2D projections of the same object.

In related work, common-line-based embeddings were learned by a neural network model to represent embeddings of 2D projections from heterogeneous samples, along with a variational autoencoder trained to reconstruct 3D objects from X-ray images [12]. Given the success of both methods, in this work, we combine representations learned from common lines via graph neural networks and learned image embeddings from pretrained and supervised CNNs to construct embeddings of 2D projections, which are then clustered into their respective objects. Different applications in computational biology have applied frameworks such as these; for example, clustering was performed on embeddings from a Siamese network trained on must-link and cannot-link sequence similarities to find groups of sequences from the same metagenome [13]. To learn fine-tuned image embeddings in our application, we trained Siamese networks with similar and dissimilar projections, learning accurate image embeddings.

In this work, for the task of clustering 2D projections of different particles into their respective groups, we present DeepSLICEM, in which we extensively explored alternative image representations and graph neural networks to improve the performance of SLICEM. Specifically, we explored 7 image representation methods with deep neural networks (VGG, ResNet-15, DenseNet, AlexNet, Efficient-Net-B1 and B7, and Siamese), 10 similarity graph node representation methods with graph neural networks (Node2Vec, Metapath2Vec, GraphWave, Watch Your Step, Attri2Vec, GraphSAGE, Deep Graph Infomax using each of the models - GCN, GAT, APPNP, and Cluster-GCN), 4 clustering methods (Birch, DBSCAN, Affinity Propagation, OPTICS) and 8 graph clustering methods (strongly and weakly connected components, walk trap, edge betweenness, greedy modularity, k-clique, semi synchronous and asynchronous label propagation).

On an experimental dataset and 5 synthetic datasets of varying clustering difficulty (obtained by adding noise to the images and changing the number of projections and classes of particles), the best methods from DeepSLICEM robustly achieve high clustering accuracy, with the pipeline yielding 92 combinations of methods across the 6 datasets, which achieve better clustering accuracy than the previous method from SLICEM (graph clustering with Walktrap or edge betweenness community detection). The improved performance underscores the merit of learning translationally and rotationally invariant image features extracted from deep neural networks and combining them with representations of features from common line-based similarity graphs using graph neural networks. We develop DeepSLICEM as an efficient AutoML pipeline for clustering 2D projections of different objects, available at https://github.com/marcottelab/2D_projection_clustering.

## Materials and methods

DeepSLICEM separates images of 2D projections of different particles into their appropriate respective groups. This is achieved by clustering the vector representations of the particles using different clustering algorithms. The SLICEM algorithm is used to construct a similarity graph of the 2D projections based on the similarity between their common lines. The final image representations are found by combining image embeddings of the images using Siamese neural networks with graph node representations of the images from the similarity graph via graph neural networks.

### Experimental datasets

We used the experimental and synthetic datasets from SLICEM [10]. The experimental dataset has a total of 100 2D projections from 4 complexes - 40S, 60S, and 80S ribosomes, and apoferritin. The synthetic dataset has a total of 204 2D projections (2-12 projections per complex) from 35 PDB entries-1A0I, 1HHO, 1NW9, 1WA5, 3JCK, 5A63, 1A36, 1HNW, 1PJR, 2FFL, 3JCR, 5GJQ, 1AON, 1I6H, 1RYP, 2MYS, 3VKH, 5VOX, 1FA0, 1JLB, 1S5L, 2NN6, 4F3T, 6B3R, 1FPY, 1MUH, 1SXJ, 2SRC, 4V6C, 6D6V, 1GFL, 1NJI, 1TAU, 3JB9 and 5A1A. To better represent real experimental conditions, we additionally construct a synthetic noisy dataset by adding random noise comprising Gaussian noise (the mean and variance of the pixels in the image are used as parameters) to simulate the grainy texture of the raw micrographs, and salt and pepper noise is added to 1% of the pixels to simulate hot and dead pixels. Two examples of noisy images compared to clean images are given in **Figure 2**.

**Figure 2.**
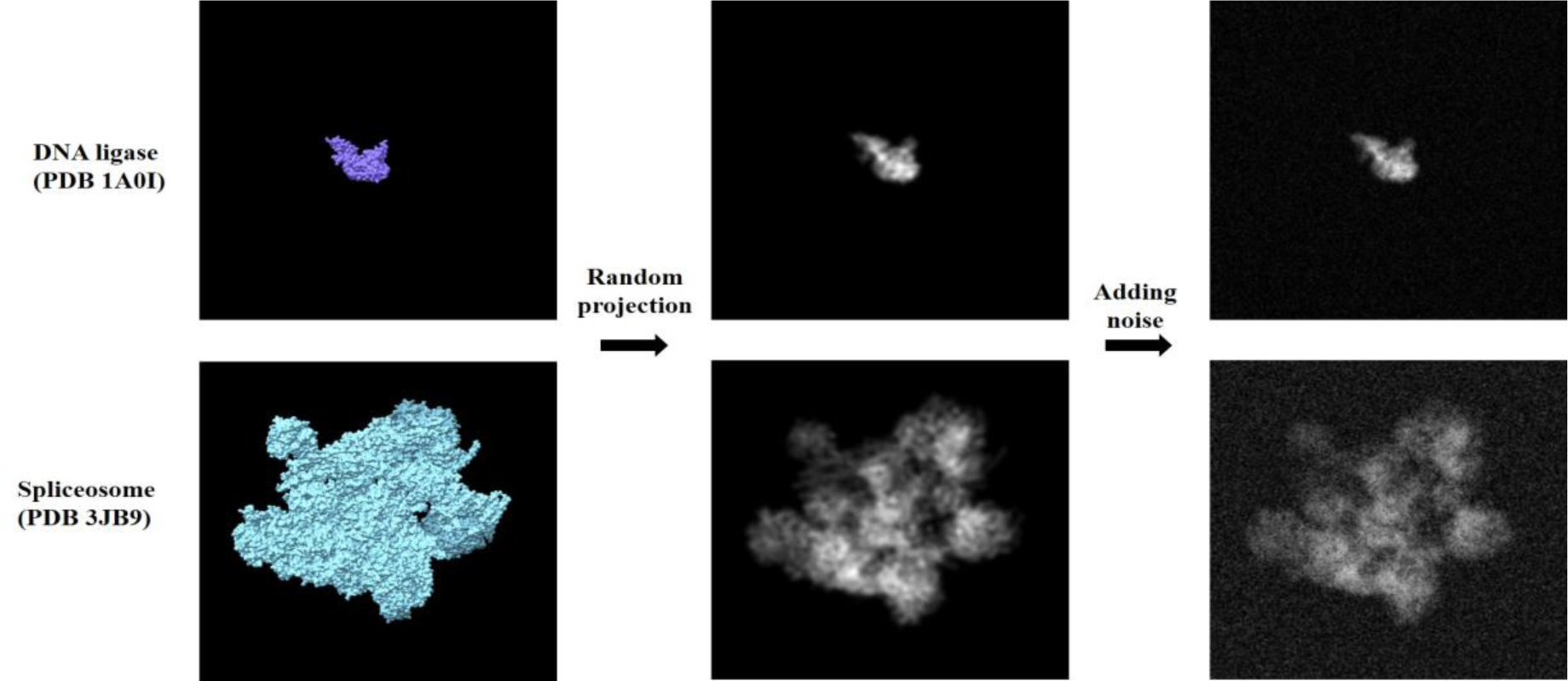
Noisy projection images are constructed from 3D molecules to build a synthetic dataset for analysis. Examples are shown here for random projections of PDB 1A0I (top) - ATP-dependent DNA ligase and 3JB9 (bottom) - Spliceosome.

We also investigated the effect of sampling more projections per complex and constructed a ‘synthetic more projections’ dataset, and its corresponding noisy dataset with a total of 558 projections (13-20 projections per complex, randomly sampled from 20-30 uniformly projected, centered 2D images of the 3D structure at a resolution of 9Å per pixel using EMAN1.9’s pdb2mrc function). All the images are rescaled to 100×100 pixels after padding the smaller images to the largest image size. To examine the potential for bias contributed by the largest complex, the ribosome, we additionally removed the ribosome to construct a ‘synthetic more projections without big ribosome’ dataset and compared this performance to study its impact. For the datasets with more projections, we resized the projections to 100 × 100 pixels from the original 350 × 350 pixels in the synthetic dataset for better memory management. To train a Siamese neural network for the experimental dataset with more samples, we also construct a combined experimental-synthetic dataset by combining the experimental dataset with the ‘synthetic more projections noisy’ dataset after resizing the 2D projections from the experimental dataset to 100 × 100 pixels from their original 96 × 96 pixels. All the images are converted into RGB format before their embeddings are found or trained in the algorithm.

### Representing projections with image embedding techniques

The 2D projection images are represented in vector space using image embeddings extracted from a convolutional neural network-based method. We explored the use of 6 different pretrained neural networks (AlexNet [14], VGG-11 [15], DenseNet [16], ResNet-18 [17], EfficientNet-B1, and EfficientNet-B7 [18]) to extract unsupervised representations of images using the img2vec python library [19].

#### Siamese neural network embedding

We also implemented a supervised learning strategy using a Siamese neural network [20], built on a ResNet-50 (with weights pretrained on ImageNet [21]) to fine-tune the embeddings such that 2D projections of the same complex are closer in vector space than those from different complexes. This is achieved by learning to estimate the similarity between images, training on triples of images comprising an anchor image (*A*), another projection of the same complex (positive image *P*), and a projection image of a different complex (negative image *N*). For this purpose, the Siamese neural network comprises 3 identical subnetworks that generate embeddings for each of the images in the triple and compare them using a triplet loss function (*L*) (adapted from the loss function of FaceNet [22]):

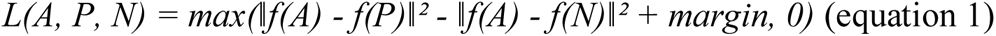

Here, *f* is the image embedding, *i.e.*, the map from an image to its representation in vector space, and *margin* is a hyperparameter that tries to enforce a margin between pairs of similar projections and dissimilar projections, which we define as 0.5 in our experiments. The two distances for the loss are returned by the custom distance layer we define in the Siamese neural network. We use a batch size of 2 for better memory management. We connect 3 dense layers (512, 256, and 256 units) to ResNet-50, with the first 2 using a ReLU activation function and batch normalization. All the weights of all the layers of the ResNet-50 model till the layer ‘conv5_block1_out’ are frozen, while the others are trainable. The model was trained using the Adam optimizer with a learning rate of 0.0001 using 10 epochs.

To find the most accurate representations of the projections trainable with the current Siamese network architecture and parameters, and to evaluate the accuracy of the subsequent clustering methods alone, we perform an experiment training the Siamese neural network on all the data. For accurate evaluation of the entire pipeline on testing data, we perform an experiment training the Siamese neural network on a training set of complexes, obtained with a 70–30 split on the complexes into training and testing data (for this experiment, we use a batch size of 32). We generate triples for each pair of 2D projections of a complex by sampling 10 negatives randomly from the set of projections of the other complexes. We also experimented by randomly sampling a negative projection image from each of the other complexes for each pair of projections from the same complex. Note that for the dataset combining both the experimental and synthetic image sets, triples are generated separately for each set and then combined. The triples dataset is split into training and validation sets using an 80–20 split. The triples dataset is split into training and validation sets using an 80–20 split. We built 7 Siamese neural networks corresponding to the 7 datasets, referred to as Siamese - synthetic, Siamese-synthetic noisy, Siamese - synthetic more projections, Siamese - synthetic more projections noisy, Siamese - more projections without ribosome, Siamese - real, and Siamese - real_synthetic. The additional Siamese neural network built using more negatives for the synthetic dataset is denoted as Siamese - synthetic more negatives.

### Incorporating projection similarities using graph node embedding techniques

The similarity graph between the 2D projections is obtained using the SLICEM algorithm [10]. Here, the similarity edge weight between two nodes (2D projection images) is the highest similarity measure out of all-by-all pairwise similarities between 1D projections of the 2D projection images projected at different angles, spanning 0 to 180 degrees at 5-degree intervals. We experiment with different similarity measures, the L1 and L2 norms. For the synthetic dataset, we also try the similarity measures - cosine similarity, correlation coefficient, and Wasserstein distance. For each node, the edge weights are updated as the Z-score relative to all edge scores, and the 5 neighbors with the highest edge weights are chosen to construct a directed similarity graph. For the experimental dataset, we also evaluate a similarity graph with the top 3000 edge weights in the original graph.

The nodes of the similarity graph are represented in vector space using a graph neural network-based node embedding method. We explore 10 different unsupervised graph node embedding methods, of which 4 methods (Node2Vec [23], Metapath2Vec [24], GraphWave [25], and Watch Your Step [26]) solely represent nodes based on the graph structure, while 6 methods (Attri2Vec [27], GraphSAGE [28], Deep Graph Infomax [29] using each of the models - GCN [30], GAT [31], APPNP [32], and Cluster-GCN [33]) incorporate image embeddings as node attributes and use this information along with the graph structure to represent nodes. All the methods were applied using the Python StellarGraph library [34].

### Combining image and graph representations of projections

We concatenated the image and graph node embeddings and applied dimensionality reduction via PCA [35] if the vectors were dense or truncated SVD [36] if they were sparse (sparsity > 0.5) and removed dimensions contributing less than 2% of the variance. We also alternatively applied dimensionality reduction to the image and graph node embeddings separately before concatenating them to yield the final embedding vectors.

### Clustering the embeddings to separate complexes

The final embeddings for the 2D projections were clustered with 4 different clustering methods (DBSCAN [37], OPTICS [38], BIRCH [39], and Affinity Propagation [40]) using the scikit-learn library [41]. We perform a 70-30 split on the complexes into training and testing data (the same split as was used for the Siamese neural network training) and choose the hyperparameters for each clustering algorithm that give the best FMMF score [42] on the training data. For the clustering algorithm, we also choose the best distance measure out of 22 measures, including linear and nonlinear metrics (Bray-Curtis, Canberra, Chebyshev, city-block, Pearson correlation, cosine similarity, Dice dissimilarity, Euclidean, Hamming, Jaccard, Jensen-Shannon, Kulsinski, Mahalanobis, matching dissimilarity, Minkowski, Rogers Tanimoto, Russell-Rao, standardized Euclidean, Sokal-Michener, Sokal-Sneath, squared Euclidean, and Yule) that gives, which yield the best silhouette score [43] on the training data. The hyperparameter ranges explored are given in **Table 1**.

**Table 1.** Hyperparameter ranges explored for the clustering algorithms.

| Method | Parameter | Range |
| --- | --- | --- |
| DBSCAN | eps | 0.25 to 3 in intervals of 0.25 |
|  | min_samples | 2 to 10 in intervals of 1 |
| OPTICS | max_eps | 0.25 to 3 in intervals of 0.25 |
|  | min_samples | 2 to 10 in intervals of 1 |
| BIRCH | threshold | 0.1 to 1 in intervals of 0.1 |
|  | branching_factor | 10 to 100 in intervals of 10 |
| AffinityPropagation | damping | 0.5 to 1 in intervals of 0.1 |

Apart from clustering the final embeddings, we also directly cluster the similarity graph (represented both as an undirected and directed graph) using different graph clustering techniques, namely, connected components, walk trap [44] (with the number of steps as 4) and edge betweenness [45], using their default parameters in the igraph library [46] and greedy modularity [47], k-clique [48], semi synchronous [49] and asynchronous [50] label propagation, using their default parameters in the networkx library [51]. The RL community detection method from [52] was also evaluated on the synthetic dataset. All the clustering methods chosen do not need an estimate of the number of clusters, making them especially suitable for this application.

### Evaluating learned complexes and their representations

The resulting clusters are compared with the ground truth clusters using different evaluation metrics, including the F-similarity-based Maximal Matching F-score (FMMF), Community-wise Maximum F-similarity-based F-score (CMMF), and Unbiased Sn-PPV Accuracy (UnSPA), which are defined in Super.Complex [42] and Qi *et al*. F-score [53], F-grand k-clique, and F-weighted k-clique [54]. We also computed unsupervised scores of the clustering using the silhouette score, Calinski-Harabasz score [55], and the Davies-Bouldin score [56]. The clustering results reported in the tables are evaluations performed using all the learned complexes compared with all known complexes. Several ‘junk’ particle images do not correspond to any complex for the experimental dataset. For this case, we report the evaluation measures, both while comparing the learned complexes with the junk particle projections and after removing the junk particle projections.

We also evaluate the quality of the embeddings for each of the projections with t-SNE [57] plots using the best distance measure, corresponding to the highest silhouette score on the training data.

## Results and Discussion

### DeepSLICEM improves on SLICEM clustering accuracy of particles in synthetic and experimental datasets

DeepSLICEM trains several clustering methods in a semi supervised fashion, as discussed in the Methods section, and selects the best method, achieving better performance on different datasets when compared to the unsupervised SLICEM method, which is currently the only other available algorithm for performing this specific task (**Table 2**). Comparing the performance across different datasets (**Table 2**, **Figures 3** and **4**), we observe that both DeepSLICEM and SLICEM achieve better performance on the (i) clean dataset than on the noisy dataset, indicating, as can be expected, that adding noise makes the clustering task harder, and (ii) on a dataset with more 2D projections than on a smaller number of projections, showing that with more 2D orientations available, more pairwise similarity data is available, making the clustering task easier. While SLICEM, like DeepSLICEM, achieves perfect performance on a clean dataset with more projections, its performance degrades more than that of DeepSLICEM on adding noise or decreasing the number of projections. This could be attributed to DeepSLICEM being more robust to translational and rotational variation than SLICEM, which is based on the theoretical property of common lines between two 2D projections of the same object. The robustness of both SLICEM and DeepSLICEM to minor variations in the number of clusters is demonstrated by continuing to achieve perfect performance when the projections corresponding to one of the clusters, the ribosome, are removed from the dataset with more synthetic projections.

**Figure 3.**
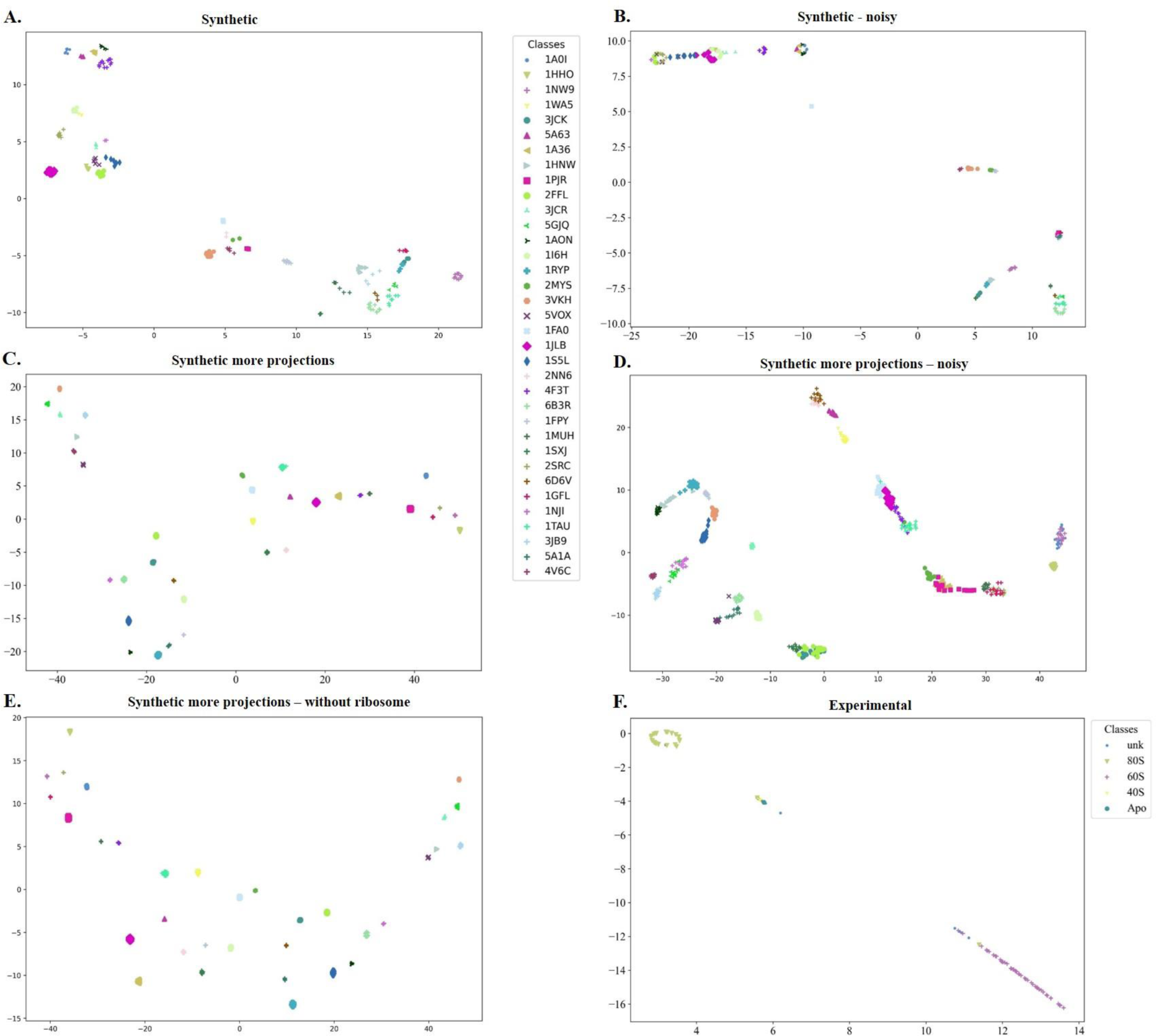
t-SNE plots of DeepSLICEM’s Siamese embeddings of different datasets show good clustering for the experimental and synthetic datasets, with the clustering improving with increasing number of projections and reducing noise (as demonstrated with synthetic data). A. Synthetic B. Synthetic - noisy C. Synthetic more projections D. Synthetic more projections noisy E. Synthetic more projections w/o ribosome F. Experimental

**Figure 4.**
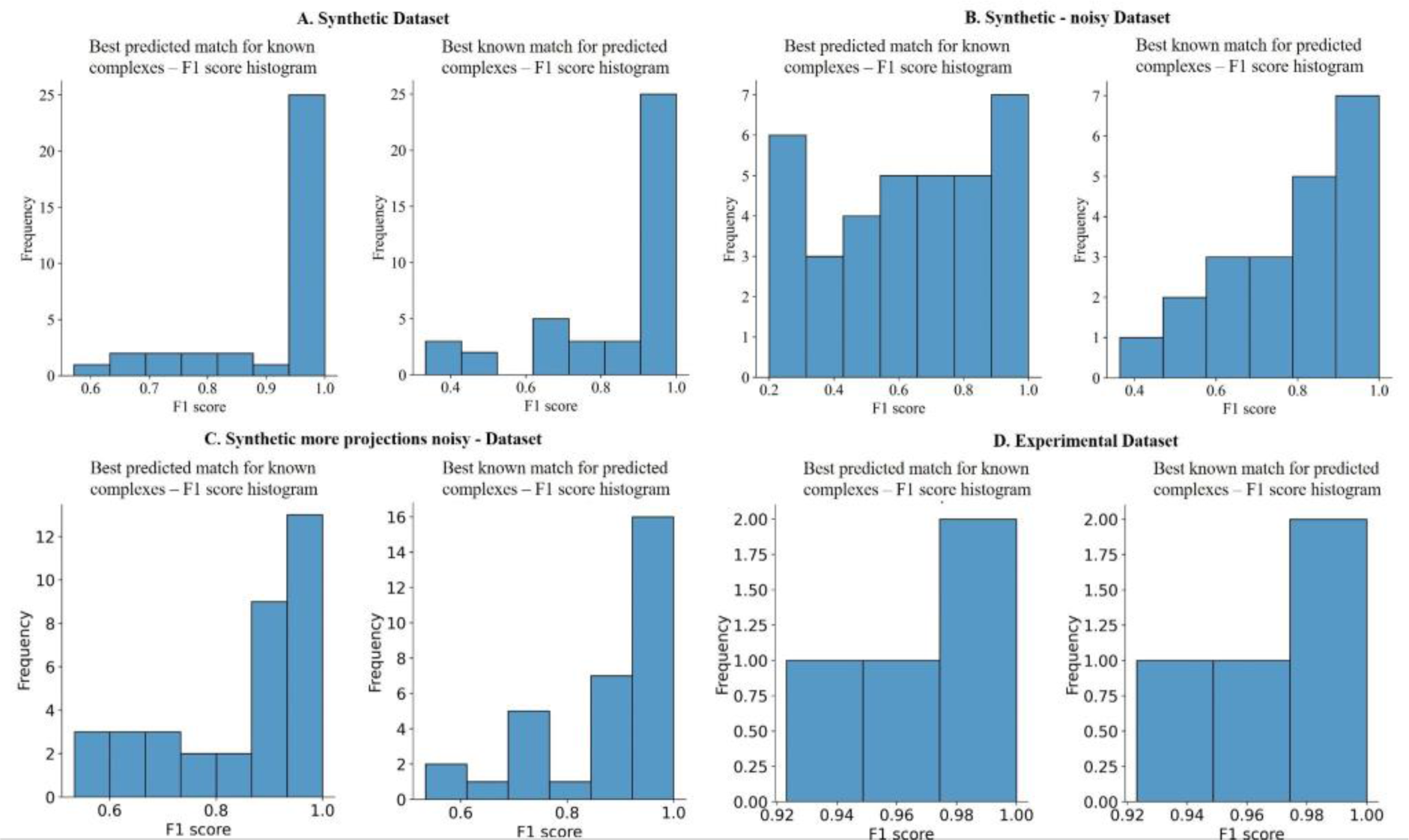
F1 score histograms of DeepSLICEM’s best clustering option on different datasets show high clustering F1 scores. A. Synthetic B. Synthetic - noisy C. Synthetic more projections noisy D. Experimental

**Table 2.**
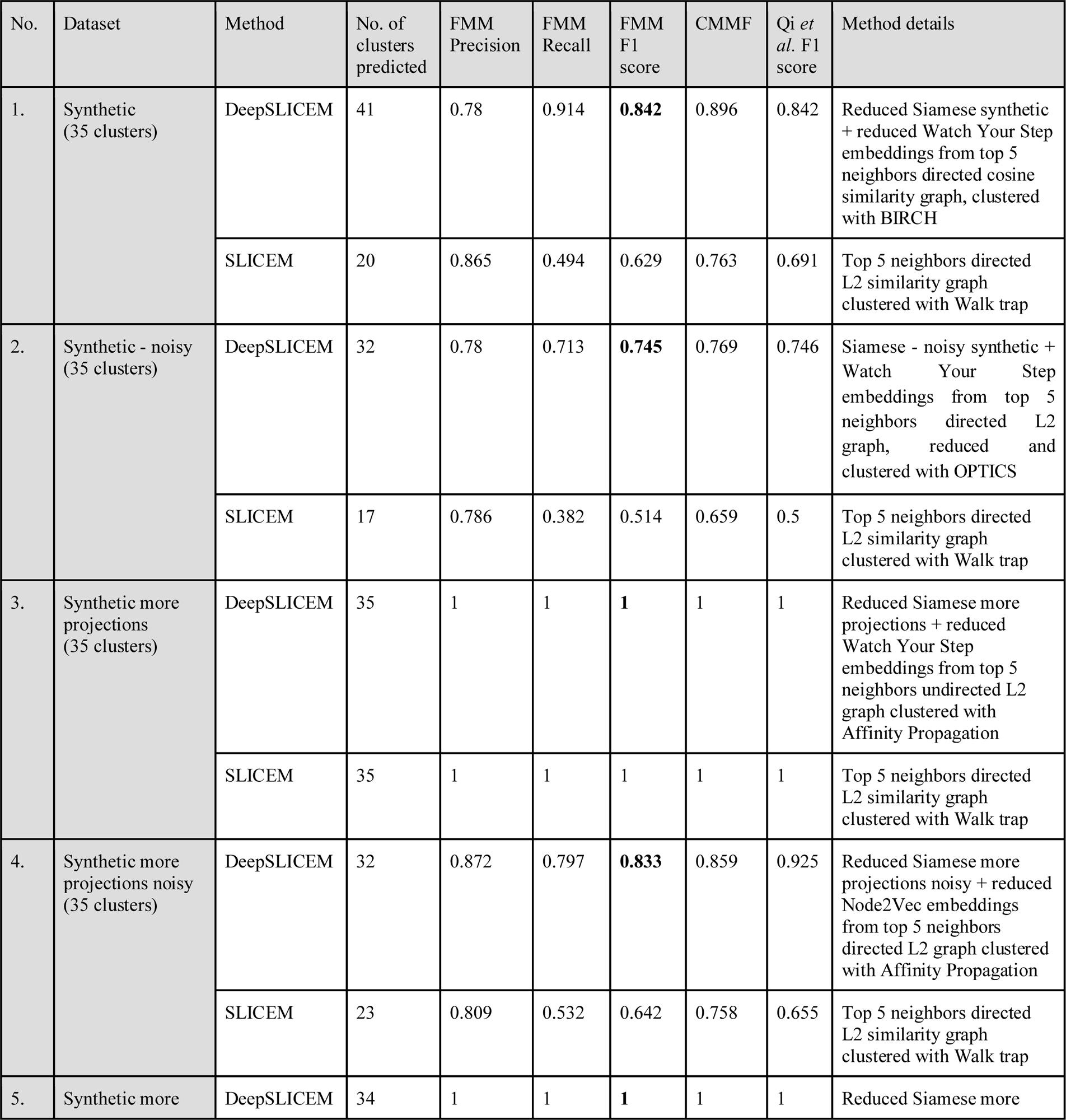
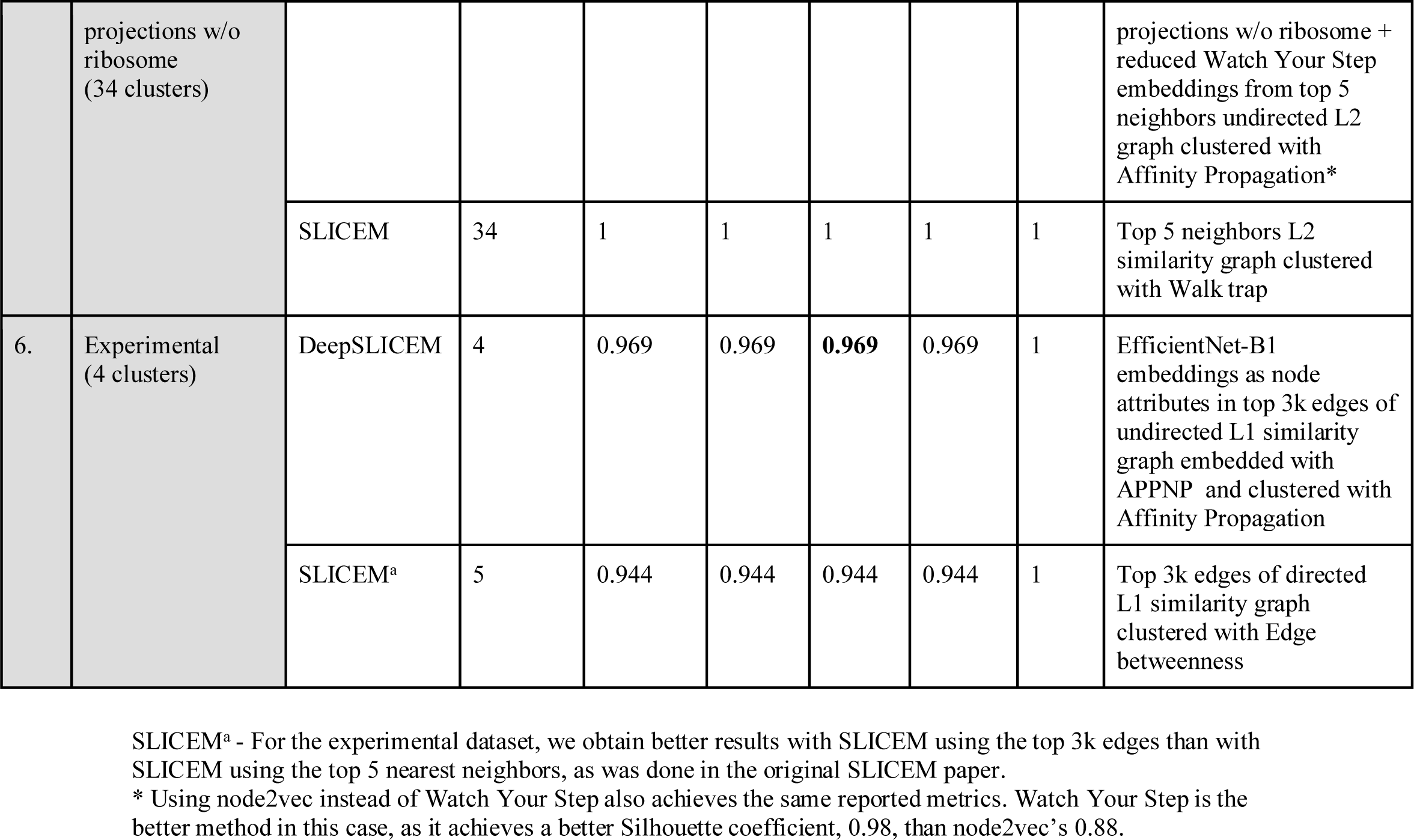
DeepSLICEM improves upon SLICEM in clustering 2D projections of different particles into their respective clusters.

| No. | Dataset | Method | No. of clusters predicted | FMM Precision | FMM Recall | FMM F1 score | CMMF | Qi *et al.* F1 score | Method details |
| --- | --- | --- | --- | --- | --- | --- | --- | --- | --- |
| 1. | Synthetic (35 clusters) | DeepSLICEM | 41 | 0.78 | 0.914 | **0.842** | 0.896 | 0.842 | Reduced Siamese synthetic + reduced Watch Your Step embeddings from top 5 neighbors directed cosine similarity graph, clustered with BIRCH |
|  |  | SLICEM | 20 | 0.865 | 0.494 | 0.629 | 0.763 | 0.691 | Top 5 neighbors directed L2 similarity graph clustered with Walk trap |
| 2. | Synthetic - noisy (35 clusters) | DeepSLICEM | 32 | 0.78 | 0.713 | **0.745** | 0.769 | 0.746 | Siamese - noisy synthetic + Watch Your Step embeddings from top 5 neighbors directed L2 graph, reduced and clustered with OPTICS |
|  |  | SLICEM | 17 | 0.786 | 0.382 | 0.514 | 0.659 | 0.5 | Top 5 neighbors directed L2 similarity graph clustered with Walk trap |
| 3. | Synthetic more projections (35 clusters) | DeepSLICEM | 35 | 1 | 1 | **1** | 1 | 1 | Reduced Siamese more projections + reduced Watch Your Step embeddings from top 5 neighbors undirected L2 graph clustered with Affinity Propagation |
|  |  | SLICEM | 35 | 1 | 1 | 1 | 1 | 1 | Top 5 neighbors directed L2 similarity graph clustered with Walk trap |
| 4. | Synthetic more projections noisy (35 clusters) | DeepSLICEM | 32 | 0.872 | 0.797 | **0.833** | 0.859 | 0.925 | Reduced Siamese more projections noisy + reduced Node2Vec embeddings from top 5 neighbors directed L2 graph clustered with Affinity Propagation |
|  |  | SLICEM | 23 | 0.809 | 0.532 | 0.642 | 0.758 | 0.655 | Top 5 neighbors directed L2 similarity graph clustered with Walk trap |
| 5. | Synthetic more projections w/o ribosome (34 clusters) | DeepSLICEM | 34 | 1 | 1 | **1** | 1 | 1 | Reduced Siamese more projections w/o ribosome + reduced Watch Your Step embeddings from top 5 neighbors undirected L2 graph clustered with Affinity Propagation* |
|  |  | SLICEM | 34 | 1 | 1 | 1 | 1 | 1 | Top 5 neighbors L2 similarity graph clustered with Walk trap |
| 6. | Experimental (4 clusters) | DeepSLICEM | 4 | 0.969 | 0.969 | **0.969** | 0.969 | 1 | EfficientNet-B1 embeddings as node attributes in top 3k edges of undirected L1 similarity graph embedded with APPNP and clustered with Affinity Propagation |
|  |  | SLICEM<sup>a</sup> | 5 | 0.944 | 0.944 | 0.944 | 0.944 | 1 | Top 3k edges of directed L1 similarity graph clustered with Edge betweenness |

SLICEM<sup>a</sup> - For the experimental dataset, we obtain better results with SLICEM using the top 3k edges than with SLICEM using the top 5 nearest neighbors, as was done in the original SLICEM paper.
\* Using node2vec instead of Watch Your Step also achieves the same reported metrics. Watch Your Step is the better method in this case, as it achieves a better Silhouette coefficient, 0.98, than node2vec's 0.88.

### Clustering combined graph node embeddings and image embeddings outperforms clustering them individually

A case study on the synthetic noisy dataset, the most difficult of the datasets evaluated, is discussed here. First, we cluster the directed similarity graph directly using different graph clustering methods (**Table 3**) and find that asynchronous label propagation performs the best (0.61 FMMF). We then explore clustering the graph node embeddings in vector space (**Table 4**) and achieve better performance than before with Watch Your Step and OPTICS clustering (0.73 FMMF). Next, we cluster image embeddings alone and observe the best performance (0.4 FMMF) with the embeddings from a Siamese neural network trained on the same dataset, which outperforms clustering other unsupervised (pre-trained) embeddings (**Table 5**), demonstrating the importance of fine-tuning a neural network to the dataset in question for better performance. We then combined the graph node embeddings with image embeddings and obtained the best performance (0.75 FMMF) clustering for Siamese - noisy synthetic embeddings concatenated with Watch Your Step, followed by dimensionality reduction and OPTICS clustering. This approach marginally improves the best results from the graph embedding clustering (0.73 FMMF), showing that the image embeddings add information to the graph node embeddings to aid the clustering.

**Table 3.** Five classical graph clustering methods were evaluated for clustering the similarity graph (constructed with 5 nearest neighbors using the L2 norm) for the synthetic noisy dataset. Asynchronous label propagation on the directed graph outperforms other methods, including walk trap on the directed graph (the algorithm used by SLICEM). For reference, the true number of clusters here is 35.

| Method | Graph type | No. of clusters | FMM Precision | FMM Recall | FMM F1 score | CMMF | Qi *et al.* F1 score |
| --- | --- | --- | --- | --- | --- | --- | --- |
| Asynchronous label propagation | Directed | 21 | 0.815 | 0.489 | **0.611** | 0.719 | 0.607 |
| Asynchronous label propagation | Undirected | 22 | 0.749 | 0.471 | 0.578 | 0.685 | 0.491 |
| Label propagation | Undirected | 20 | 0.733 | 0.419 | 0.533 | 0.645 | 0.473 |
| Walktrap (SLICEM) | Directed | 17 | 0.786 | 0.382 | **0.514** | 0.659 | 0.500 |
| Greedy modularity | Directed | 14 | 0.788 | 0.315 | 0.451 | 0.619 | 0.408 |
| Greedy modularity | Undirected | 13 | 0.767 | 0.285 | 0.416 | 0.588 | 0.375 |
| k-clique | Undirected | 7 | 0.577 | 0.115 | 0.192 | 0.399 | 0.048 |

**Table 4.** Four graph neural network based clustering methods were evaluated for clustering the (directed) similarity graph (constructed with 5 nearest neighbors using the L2 norm) for the synthetic noisy dataset. Watch your step node embeddings with OPTICS clustering outperforms other methods. For reference, the true number of clusters here is 35. Note that only the best clustering method (out of 4 clustering methods) for each of the embeddings is reported in this table.

| Node Embedding Method | Clustering Method | Clustering parameters | No. of clusters | FMM Precision | FMM Recall | FMM F1 score | CMMF | Qi *et al.* F1 score |
| --- | --- | --- | --- | --- | --- | --- | --- | --- |
| Watch Your Step | OPTICS | max_eps=2.75, metric='cosine', min_samples=3 | 31 | 0.777 | 0.688 | **0.73** | 0.773 | 0.788 |
| Node2Vec | OPTICS | max_eps=1.5, metric='cosine' | 22 | 0.776 | 0.488 | **0.6** | 0.698 | 0.632 |
| Metapath2Vec | OPTICS | max_eps=1.5, metric='cosine' | 18 | 0.744 | 0.383 | 0.505 | 0.63 | 0.491 |
| GraphWave | Affinity Propagation | damping=0.9 | 17 | 0.408 | 0.198 | 0.267 | 0.353 | 0.077 |

**Table 5.** Seven CNN-based image embedding methods were evaluated in terms of clustering performance on synthetic noisy dataset images. The ResNet-50-based Siamese network trained on noisy synthetic data outperforms other methods by a significant margin, demonstrating the improvement achieved by fine-tuning embeddings to the dataset being evaluated. Note that only the best clustering method (out of 4 clustering methods) for each of the embeddings is reported in this table.

| Embedding Method | Clustering Method | Clustering parameters | FMM Precision | FMM Recall | FMM F1 score | CMMF |
| --- | --- | --- | --- | --- | --- | --- |
| Siamese - noisy synthetic | BIRCH | branching_factor=90<br>threshold=0.2 | 0.576 | 0.313 | **0.405** | 0.511 |
| Siamese - synthetic | BIRCH | branching_factor=20<br>threshold=0.4 | 0.088 | 0.280 | 0.134 | 0.136 |
| ResNet-18 | BIRCH | branching_factor=10<br>threshold=0.9 | 0.077 | 0.322 | 0.124 | 0.124 |
| Siamese - synthetic more projections | BIRCH | branching_factor=10<br>threshold=0.1 | 0.081 | 0.244 | 0.121 | 0.125 |
| EfficientNet-B1 | BIRCH | branching_factor=90<br>threshold=0.1 | 0.074 | 0.315 | 0.120 | 0.120 |
| DenseNet | BIRCH | branching_factor=10<br>threshold=0.9 | 0.066 | 0.348 | 0.111 | 0.111 |
| AlexNet | Affinity Propagation | damping=0.5 | 0.121 | 0.055 | 0.076 | 0.112 |
| EfficientNet-B7 | Affinity Propagation | damping=0.9 | 0.110 | 0.057 | 0.075 | 0.109 |
| VGG | Affinity Propagation | damping=0.6 | 0.112 | 0.054 | 0.073 | 0.110 |

### Multiple methods in DeepSLICEM achieve better clustering than SLICEM’s Walktrap graph clustering

For this task of image clustering with the help of an image similarity graph, DeepSLICEM finds 92 method combinations (from 7 CNN-based image representation methods, 10 GNN-based graph node representation methods, 4 spatial clustering methods, and 8 graph clustering methods) that achieve better clustering accuracy than previous methods from SLICEM on 7 cryo-EM projection image datasets. The best methods chosen by DeepSLICEM include ResNet-50 Siamese neural network-based image embeddings, Watch Your Step graph node embeddings, OPTICS (spatial) clustering, and asynchronous label propagation graph clustering. In this section, we report the methods from DeepSLICEM that outperform SLICEM for each type of dataset, thus reporting the best methods for clean data, noisy data, limited data, and extended data. Next, we analyze each method for its ability to outperform SLICEM in all the experiments to benchmark different methods and recommend the best methods for a new dataset to improve the runtime of an AutoML approach by exploring several method combinations.

We first report the performance of DeepSLICEM on a noisy dataset based on its closer resemblance to an experimental dataset when compared to the other types of synthetic datasets constructed. While SLICEM (walk trap clustering on L2 similarity graph) achieves 0.51 FMMF, 30 methods attempted by DeepSLICEM yield better performance. These methods include 3 graph clustering methods - asynchronous and synchronous label propagation on the directed graph, and synchronous label propagation on the undirected graph (**Table 3**); 2 methods clustering graph node embeddings in vector space - Watch Your Step and Node2Vec embeddings clustered by the OPTICS algorithm (**Table 4**); 7 methods clustering reduced dimension vectors from image embeddings concatenated with graph node embeddings (**Table 6**); 13 methods clustering reduced image embeddings concatenated with reduced graph node embeddings (**Table S1**); and 5 methods clustering graph node embeddings with image embeddings incorporated as node attributes (**Table S2**).

**Table 6.** Combined image embeddings and graph node embeddings were evaluated in terms of clustering performance on synthetic noisy dataset images. The results presented in this table are all the DeepSLICEM methods (clustering combined image and graph node embeddings), which yield better performance than SLICEM. The best performance for DeepSLICEM was obtained with image embeddings from the Siamese network trained on noisy synthetic data, concatenated with Watch Your Step graph node embeddings, and clustered with OPTICS.

| Image embedding Method | Node Embedding Method | Clustering Method | Clustering parameters | No. of clusters | FMM Precision | FMM Recall | FMM F1 score | CMMF | Qi *et al.* F1 score |
| --- | --- | --- | --- | --- | --- | --- | --- | --- | --- |
| Siamese - noisy synthetic | Watch Your Step | OPTICS | max_eps=0.5, metric='cosine', min_samples=3 | 32 | 0.780 | 0.713 | **0.745** | 0.769 | 0.746 |
| Siamese - synthetic more projections | Watch Your Step | OPTICS | max_eps=2.75, metric='cosine', min_samples=3 | 32 | 0.766 | 0.700 | 0.731 | 0.766 | 0.746 |
| Siamese - noisy synthetic | Node2Vec | DBSCAN | metric='sqeuclidean', min_samples=3 | 22 | 0.857 | 0.539 | 0.662 | 0.762 | 0.667 |
| Siamese - synthetic more projections | Node2Vec | OPTICS | max_eps=0.25, metric='cosine', min_samples=4 | 23 | 0.771 | 0.507 | 0.612 | 0.705 | 0.655 |
| Siamese - synthetic | Watch Your Step | OPTICS | max_eps=0.5, metric='cosine' | 21 | 0.788 | 0.473 | 0.591 | 0.696 | 0.643 |
| Siamese - noisy synthetic | GraphWave | OPTICS | max_eps=2.25, metric='sqeuclidean', min_samples=3 | 30 | 0.605 | 0.518 | 0.558 | 0.613 | 0.462 |
| Siamese - synthetic | Node2Vec | Affinity Propagation | damping=0.9 | 19 | 0.788 | 0.428 | 0.555 | 0.679 | 0.519 |

Next, we discuss the effect of improving the noisy synthetic dataset by adding more projections of the images, corresponding to an experimental scenario where more data on different projections can be collected. On this ‘synthetic more projections noisy dataset’, compared to SLICEM (walk trap clustering on L2 similarity graph), which achieves 0.64 FMMF, 31 methods attempted by DeepSLICEM achieve better performance, *i.e.*, 3 graph clustering methods - on the undirected graph, asynchronous label propagation (achieving the best performance with 0.667 FMMF) and synchronous label propagation, and synchronous label propagation on the directed graph (**Table S3**); 13 methods clustering reduced dimension vectors from image embeddings concatenated with graph node embeddings (**Table S3**) (6 on the directed graph and 7 on the undirected graph); and 15 methods clustering reduced image embeddings concatenated with reduced graph node embeddings (**Table S3**) (8 on the directed graph and 7 on the undirected graph).

We now turn to the effect of denoising experimental datasets by considering the clean synthetic dataset. On the synthetic dataset, compared to SLICEM (walk trap clustering on L2 similarity graph), which achieves 0.63 FMMF, 27 methods attempted by DeepSLICEM yield better performance, *i.e.*, 5 graph clustering methods - asynchronous label propagation on the directed cosine similarity graph (achieving the best performance with 0.83 FMMF) and L2 similarity graph and undirected L2 similarity graph and synchronous label propagation on the undirected cosine similarity graph (**Table S4**); 4 methods clustering reduced dimension vectors from image embeddings concatenated with graph node embeddings (**Table S4**); 10 methods clustering reduced image embeddings concatenated with reduced graph node embeddings (**Table S4**); and 8 methods clustering graph node embeddings with image embeddings incorporated as node attributes (**Table S4**).

Finally, we analyze the effects of both denoising and collecting additional data on different projections. With this ‘synthetic-more-projections dataset’, one of the methods attempted by DeepSLICEM (reduced Siamese more projections + reduced Watch Your Step embeddings from the top 5 neighbors undirected L2 graph clustered with Affinity Propagation), reported in **Table 2**, outperforms SLICEM, while two methods from DeepSLICEM (reduced Siamese more projections w/o ribosome + reduced Watch Your Step or Node2vec embeddings from the top 5 neighbors undirected L2 graph clustered with Affinity Propagation), reported in **Table 2**, achieve better performance than SLICEM on the synthetic more projections dataset without the ribosome. On the experimental dataset, with the top 3k edges of the undirected L1 similarity graph, SLICEM (walk trap clustering) is outperformed by one method attempted by DeepSLICEM (**Table 2**) EfficientNet-B1 embeddings as node attributes in the graph embedded with APPNP, clustered with Affinity Propagation.

Given the plethora of methods explored by DeepSLICEM, we next investigated which methods achieved good performance consistently across datasets out of the 92 experiments in which DeepSLICEM achieved better performance than SLICEM. Among the 81 clustering vector representations, DeepSLICEM found that the best clustering method was OPTICS, followed by Affinity Propagation, Birch, and DBSCAN (**Figure 5A**). Next, we examined which node embedding methods performed well. Of the 81 experiments in which node embeddings were clustered, Watch Your Step was chosen as the best method 49% of the time, followed by node2vec (28%) and other methods, each constituting <6% of the wins (**Figure 5B**). We then examined which image embedding methods were chosen as the best (in conjunction with a node embedding method) in 74 experiments outperforming SLICEM. We observed that Siamese embeddings won most of the time compared to unsupervised methods, of which EfficientNets won most of the time (**Figure 5C**). From 11 experiments of different graph clustering methods, we find that label propagation methods on directed and undirected graphs perform well (**Figure 5D**) with asynchronous label propagation on both types of graphs dominating accuracy, as was observed earlier. From **Table 2**, we observe that the best-performing methods for different datasets comprise predominantly Siamese embeddings, followed by EfficientNet-B1 embeddings for image representation; Watch your step, followed by node2vec for graph node embeddings; and Affinity Propagation, followed by OPTICS and Birch for clustering - consistent with their dominance in the 92 combinations of methods performing better than SLICEM.

**Figure 5.**
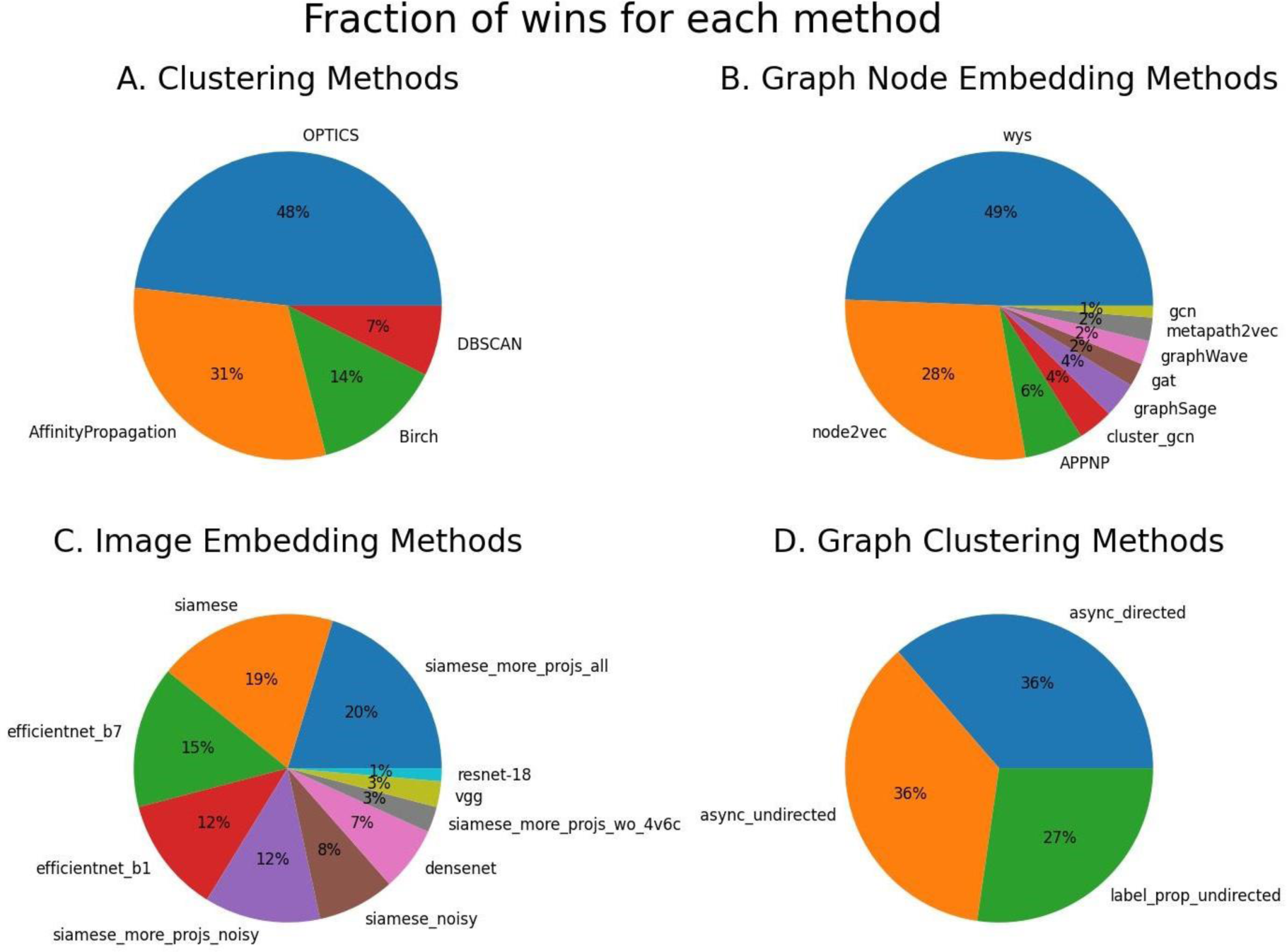
The best methods chosen by DeepSLICEM for the task of image clustering with the help of graph representation, make use of OPTICS clustering, Watch Your Step graph node embeddings, (ResNet-50) Siamese neural network-based image embeddings, and Asynchronous Label propagation. A. Clustering methods B. Graph node embedding methods C. Image embedding methods D. Graph clustering methods.

### Conclusions and future work

In this paper, we demonstrate that there is a valid path forward for using deep neural networks to accurately separate mixtures of 2D projections of different macromolecules computationally. Such an approach should help reduce the need for experimental purification of individual complexes, opening cryo-EM structure determination to considerably more heterogeneous samples. Previous work, SLICEM, has explored the use of similarity between common lines of projections to construct a pairwise similarity graph, which was then clustered using 2 methods. In this work, we develop DeepSLICEM, a pipeline that evaluates 10 graph neural networks for representing the common line similarity graph of images, combined with 7 different CNNs that add additional features from the images, and clusters the images via 13 different methods. Across 6 datasets from synthetic and experimental macromolecules, DeepSLICEM finds 91 methods (combinations of image embedding methods, graph node embedding methods, and clustering methods) that achieve better clustering accuracy than SLICEM. The best methods that dominate in terms of performance in DeepSLICEM include combinations of Watch your step graph node embeddings, clustering with Affinity Propagation, and Siamese neural network image embeddings; the latter demonstrating how supervision can help improve the performance achieved with the unsupervised method, SLICEM.

Different future directions can be explored to improve performance. The current image embeddings used are from CNNs pretrained on images from ImageNet. Since ImageNet has millions of images, on average, having ∼1000 images per synonym set, with objects represented diversely, *i.e.*, in different poses, from different viewpoints, having background noise, with occlusions, and localized to different parts of the images, the CNNs learn rotational and translational invariance of the object in question, making it suitable for the current problem of learning similar embeddings for different 2D projections of the same 3D object. However, ImageNet does not have any images of micrographs, so the weights of the CNNs can be retrained to learn Cryo-EM image features, as demonstrated with the Siamese neural network retraining a ResNet-50. This can be attempted with other CNNs that perform well, such as the EfficientNet-B1. Alternately, autoencoders such as CryoDRGN [58] can be trained on large datasets of molecules from CryoEM images to learn good image embeddings that can be used for clustering. Perspective transformer nets [59] have been proposed for transforming 2D projections of objects into 3D representations. The encoder part of their neural network can be fine-tuned for our problem and used to generate image embeddings.

Considering the success of DeepSLICEM in clustering projections of macromolecules from Cryo-EM, this AutoML pipeline can also be explored for application to any problem where different perspectives of multiple objects are to be grouped.

## Code and data availability

The software developed in this work is publicly available on GitHub at https://github.com/marcottelab/2D_projection_clustering. All the data and results files are available on Zenodo at https://doi.org/10.5281/zenodo.10614536.

## Acknowledgments

The authors are grateful to Eric Verbeke for helpful discussion and assistance with SLICEM and Caitie McCafferty for her valuable insights, assistance with EMAN, and contribution of the code for expanding the synthetic dataset. We also thank Kalyani Palukuri for making **Figure 1** of the paper. This research was funded by grants to E.M.M. from the National Institute of General Medical Sciences (R35 GM122480), the National Institute of Child Health and Human Development (R01 HD085901), and the Welch Foundation (F-1515).

