## Supplemental Tables S1 and S2 for "DeepSLICEM: Clustering CryoEM particles using deep image and similarity graph representations"

### Supplementary tables

**Table S1. Different (dimensionally) reduced image embedding methods concatenated with reduced graph node embedding methods for clustering synthetic noisy images are evaluated.** The results presented are the DeepSLICEM methods giving better performance than SLICEM.

| Image embedding Method | Node Embedding Method | Clustering Method | Clustering parameters | No. of clusters | FMM Precision | FMM Recall | FMM F1 score | CMMF | Qi *et al.* F1 score |
|---|---|---|---|---|---|---|---|---|---|
| Siamese - noisy synthetic | Watch Your Step | OPTICS | max_eps=0.25, metric='cosine', min_samples=3 | 31 | 0.79 | 0.7 | **0.742** | 0.77 | 0.758 |
| Siamese - synthetic more projections | Watch Your Step | Birch | n_clusters=None, branching_factor =50, threshold=0.8 | 27 | 0.759 | 0.585 | 0.661 | 0.748 | 0.710 |
| Siamese - noisy synthetic | Node2Vec | OPTICS | max_eps=2.75, metric='sqeuclidean', min_samples=3 | 34 | 0.639 | 0.621 | 0.630 | 0.711 | 0.609 |
| Siamese - synthetic more projections | Node2Vec | OPTICS | max_eps=0.25, metric='cosine' | 22 | 0.773 | 0.486 | 0.596 | 0.697 | 0.596 |
| EfficientNet-B7 | Watch Your Step | Affinity Propagation | damping=0.9 | 22 | 0.755 | 0.474 | 0.583 | 0.693 | 0.456 |
| VGG | Watch Your Step | OPTICS | max_eps=2.5, metric='braycurtis', min_samples=3 | 22 | 0.747 | 0.469 | 0.576 | 0.646 | 0.526 |
| EfficientNet-B7 | Node2Vec | Affinity Propagation | damping=0.9 | 23 | 0.718 | 0.472 | 0.569 | 0.648 | 0.552 |
| Siamese - noisy synthetic | Metapath2Vec | BIRCH | branching_factor =30, n_clusters=None | 19 | 0.804 | 0.437 | 0.566 | 0.691 | 0.593 |
| Siamese - synthetic | Node2Vec | Affinity Propagation | damping=0.9 | 19 | 0.796 | 0.432 | 0.560 | 0.687 | 0.556 |
| VGG | Node2Vec | Affinity Propagation | damping=0.9 | 23 | 0.674 | 0.443 | 0.534 | 0.635 | 0.483 |
| DenseNet | Watch Your Step | OPTICS | max_eps=2.5, metric='braycurtis', min_samples=4 | 20 | 0.732 | 0.419 | 0.533 | 0.629 | 0.582 |
| Siamese - synthetic | Watch Your Step | OPTICS | max_eps=0.25, metric='correlation', | 24 | 0.649 | 0.445 | 0.528 | 0.630 | 0.508 |

| | | | | | | | | | |
|---|---|---|---|---|---|---|---|---|---|
| | | | min_samples=3 | | | | | | |
| EfficientNet-B1 | Watch Your Step | OPTICS | max_eps=0.5, metric='braycurtis', min_samples=4 | 19 | 0.742 | 0.403 | 0.522 | 0.635 | 0.481 |

**Table S2. Different methods of clustering graph node embeddings using image embeddings as node attributes are evaluated for synthetic noisy images.** The results presented are the DeepSLICEM methods giving better performance than SLICEM.

| Image embedding Method | Node Embedding Method | Clustering Method | Clustering parameters | No. of clusters | FMM Precision | FMM Recall | FMM F1 score | CMMF | Qi *et al.* F1 score |
|---|---|---|---|---|---|---|---|---|---|
| EfficientNet-B7 | APPNP | BIRCH | branching_factor=80, threshold=0.3 | 34 | 0.602 | 0.584 | **0.593** | 0.674 | 0.493 |
| EfficientNet-B7 | Cluster-GCN | BIRCH | branching_factor=80 | 34 | 0.579 | 0.563 | 0.571 | 0.637 | 0.406 |
| EfficientNet-B1 | Cluster-GCN | BIRCH | branching_factor=20 | 27 | 0.622 | 0.480 | 0.542 | 0.639 | 0.484 |
| EfficientNet-B7 | GraphSage | BIRCH | branching_factor=70, threshold=0.2 | 39 | 0.505 | 0.563 | 0.532 | 0.614 | 0.351 |
| EfficientNet-B1 | APPNP | BIRCH | branching_factor=60, n_clusters=None, threshold=0.2 | 55 | 0.433 | 0.681 | 0.529 | 0.617 | 0.489 |
